# Cell surface composition and ionic strength mediate fast sedimentation in the cyanobacterium *Synechococcus elongatus* PCC 7942

**DOI:** 10.1101/2022.12.21.521370

**Authors:** Julie A. Z. Zedler, Marlene Michel, Georg Pohnert, David A. Russo

## Abstract

Cyanobacteria are photosynthetic prokaryotes of high ecological and biotechnological relevance that have been cultivated in laboratories around the world for more than 70 years. Prolonged laboratory culturing has led to multiple microevolutionary events and the appearance of a large number of “domesticated” substrains among model cyanobacteria. Despite its widespread occurrence, strain domestication is still largely ignored. In this work we describe *Synechococcus elongatus* PCC 7942–KU, a novel domesticated substrain of the model cyanobacterium *Synechococcus elongatus* PCC 7942, which presents a fast-sedimenting phenotype. Under higher ionic strengths the sedimentation rate increases leading to complete sedimentation in just 12 h. Through whole genome sequencing and gene deletion, we demonstrate that the Group 3 alternative sigma factor F (SigF) plays a key role in cell sedimentation. In addition, sedimentation analysis of an unpiliated mutant and differences in surface hydrophobicity suggest that mutations in SigF lead to significant changes of cell surface structures and, consequentially, to the appearance of a fast-sedimenting phenotype. This work sheds light on the determinants of the planktonic to benthic transitions and provides genetic targets to generate fast-sedimenting strains that could unlock cost-effective cyanobacterial harvesting at scale.

## 1. Introduction

Every axenic laboratory culture ultimately derives from a “wild” environmental ancestor. The transition from the environment to the laboratory typically implies moving from a highly fluctuating, competitive environment to a single species, static culture. The adaptation of environmental isolates to their laboratory habitat is termed domestication (Eydallin *et al*., 2014). Virtually all modern model organisms have undergone decades of domestication and cyanobacteria, globally important phototrophic prokaryotes, are not an exception. Due to the interest in their ecology, physiology and, more recently, biotechnological applications, cyanobacterial model strains have been maintained in laboratories around the world for more than 70 years (Bryant, 2014). The continuous culturing, passaging and mutagenising of cyanobacterial strains has led to domestication and the concomitant appearance of multiple substrains. The extent of which was first highlighted in the widely used *Synechocystis* sp. PCC 6803 (hereafter *Synechocystis*) (Trautmann *et al*., 2012). *Synechocystis* was originally isolated in 1968 and soon after diverged into two lineages, PCC 6803 and ATCC 27184 (Ikeuchi and Tabata, 2001). ATCC 27184 would later be used to generate the first glucose-tolerant (GT) strains (Williams, 1988). As a result of a sequenced genome (Kaneko *et al*., 1996), and their ease of cultivation (e.g. growth on glucose, lack of motility and aggregation), GT *Synechocystis* strains rapidly became popular hosts to study the genetics of cyanobacteria and, later, for a myriad of biotechnological applications. On the other hand, strains of the PCC 6803 lineage have played a key role in unravelling the mechanisms behind motility, phototaxis and protein secretion (Bhaya *et al*., 1999; Schuergers *et al*., 2016; Russo *et al*., 2022).

A similar pattern of laboratory domestication is now emerging for another well-studied freshwater cyanobacterium, *Synechococcus elongatus* (hereafter *S. elongatus*) (Adomako *et al*., 2022). *S. elongatus* PCC 7942 has been used extensively as a model for the prokaryotic circadian clock (Cohen and Golden, 2015). Recently, it has also been used to study natural competence (Schirmacher *et al*., 2020; Taton *et al*., 2020) and developed as a biotechnological chassis (Santos-Merino *et al*., 2019). The origin of *S. elongatus* can be tracked back to early isolations in Waller Creek, Texas and later in San Francisco, California (Stanier *et al*., 1971; Golden, 2019). Several *S. elongatus* strains were isolated and deposited in the Pasteur Culture Collection, however only PCC 6301 (alias UTEX 625) and PCC 7942 were widely adopted. In 2015, a fast-growing strain of *S. elongatus*, UTEX 2973, was isolated from a frozen archive of UTEX 625 (Yu *et al*., 2015). While the exact origin of UTEX 2973 remains a mystery, it is likely that the specific growth conditions used in this laboratory (e.g. high light, high CO_2_) might have favoured the appearance of a fast-growing substrain. Another addition to the *S. elongatus* group of substrains was UTEX 3055, an environmental isolate from Waller Creek whose genome shares 98.5% identity to that of PCC 7942 (Yang *et al*., 2018).

Further evidence for laboratory domestication arises from the distinct phenotypes presented by the *S. elongatus* substrains. For example, the laboratory strains PCC 6301, PCC 7942 and UTEX 2973 share a genome identity of 99.99%. However, when compared to PCC 7942, both PCC 6301 and UTEX 2973 lack natural competence, the ability to take up and incorporate foreign DNA (Schirmacher *et al*., 2020; Taton *et al*., 2020). It has been suggested that this is due to the presence of single nucleotide polymorphisms (SNPs) in critical subunits of the type IVa pili system (T4aPS) (Li *et al*., 2018). UTEX 2973 also exhibits a fast-growing and high-light tolerant phenotype (Ungerer, Lin, *et al*., 2018; Ungerer, Wendt, *et al*., 2018). A more dramatic phenotypical difference can be seen in the environmental isolate UTEX 3055. In laboratory conditions, UTEX 3055 forms biofilms and is capable of phototactic motility with a unique bidirectional photoreceptor (Yang *et al*., 2018). PCC 7942 is not phototactic, however the genes encoding for this photoreceptor, and other components of the phototaxis pathway, are present in its genome. In addition, PCC 7942 does not form biofilms, however mutations in key T4aPS components (e.g. PilB1) lead to a strong biofilm-forming phenotype (Schatz *et al*., 2013). Therefore, it has been suggested that phototactic motility and biofilm formation are intrinsic properties of *S. elongatus* and were lost during laboratory domestication (Adomako *et al*., 2022). Interestingly, genotype-to-phenotype analyses of domesticated *Synechocystis* and *S. elongatus* strains indicate similar loss-of-function patterns. These include extensively modified cell surfaces (e.g. loss of pili, changes in polysaccharide composition) and loss of environmentally relevant behaviours such as motility, phototaxis and biofilm formation. Despite ample evidence of substrain individuality, *Synechocystis* and *S. elongatus* substrains are widely used and strain domestication has received little attention.

In this study we describe a new *S. elongatus* substrain, PCC 7942–KU (hereafter 7942– KU), that shows a fast-sedimenting phenotype. We demonstrate that fast sedimentation occurs due to changes in cell surface composition mediated, in part, by a mutation in the group 3 sigma factor, SigF. Our findings present a unique opportunity to investigate the mechanisms that underly the switch from a planktonic to a benthic state. In addition, fast sedimentation also enables efficient and low-cost harvesting for cyanobacterial bioproduction.

## 2. Results

### 7942–KU exhibits a strong sedimentation phenotype in high-density medium

7942–KU was revived from a *S. elongatus* cryo archive at the University of Copenhagen. Although the complete strain history is unknown, 7942–KU has its origins in a *S. elongatus* strain donated by Susan Golden (UC San Diego) to a European cyanobacterial research group before 2012. Given it had spent more than a decade being cultivated and transferred amongst European laboratories, we hypothesised that it could have undergone laboratory domestication. Therefore, we purchased *S. elongatus* PCC 7942 from the Pasteur Culture Collection (hereafter 7942–WT) and initiated a phenotypic comparison between both strains. It was immediately evident that 7942–KU exhibited a sedimenting phenotype when cultivated in the high-density medium P4 (Figure 1A). To further investigate this phenomenon, both strains were cultivated in BG-11 and P4 media and the sedimentation rates were investigated over 24 hours (Figure 1B). In BG-11 medium, 7942–WT does not sediment and only minor sedimentation is observed in 7942–KU. In P4 medium, three hours are sufficient for a large part of the 7942–KU culture to sediment. Full sedimentation was achieved after 12 hours with little difference observed between 12 and 24 hours. 7942–WT also exhibits a faster sedimentation rate when grown in P4 medium, however the culture never completely sedimented during the experiment. After 24 hours of static incubation, the sedimented cultures could be completely resuspended by shortly vortexing them (Figure 1C). This is distinct from the biofilm-forming phenotype previously observed in 7942–WT and UTEX 3055 where the cells strongly adhere to the culture vessel (Schatz *et al*., 2013; Yang *et al*., 2018).

**Figure 1.**
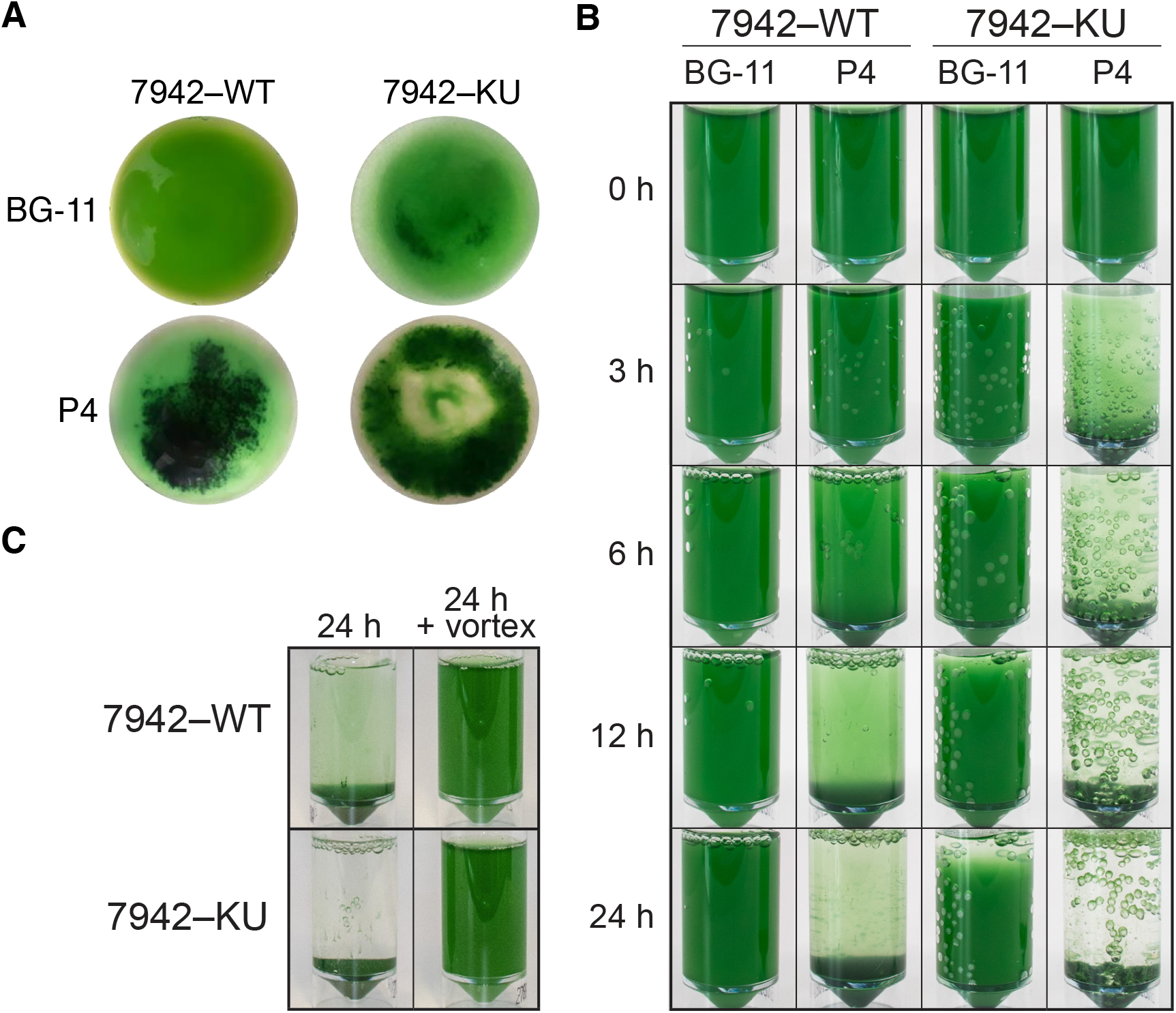
Sedimentation of 7942–WT and 7942–KU cultivated in BG-11 or P4 medium. **A** 20 mL cultures of 7942–WT and 7942–KU after a 2-week incubation in round bottom flasks under static conditions. Pictures represent a top view of the flask. **B** 24-hour sedimentation assay (15 mL) in universal containers (30 mL) incubated under static conditions. **C** 7942–WT and 7942–KU before and after 5 s of vortexing. Cultures were grown in P4 medium and incubated statically for 24 h.

### Sedimentation is triggered by an increase in ionic strength of the medium

To determine the factors that could be causing the accelerated sedimentation we started by comparing the components in BG-11 and P4 media (Supporting Information 1, Table S3). One difference that stood out was that the concentration of MgSO_4_ being almost seven times higher in P4 than in BG-11. MgSO_4_ is a popular flocculant in several industries and it has previously been used for flocculation of algal cultures at high pH (Vandamme *et al*., 2012). To test the effect of MgSO_4_ on sedimentation rates, 7942–KU was transferred to BG-11 medium with varying concentrations of the salt (0.3, 2, 4 and 8 mM) and then incubated in static conditions for 24 hours (Figure 2). Sedimentation increased with increasing MgSO_4_ concentrations with full sedimentation being observed at 8 mM. This confirmed that MgSO_4_ was directly influencing the sedimentation of the cultures, but the question remained whether this was due to a specific effect of the magnesium salt or simply due to an increase in salt concentration. To answer this question, sedimentation rates were compared with the addition of other salts.

**Figure 2.**
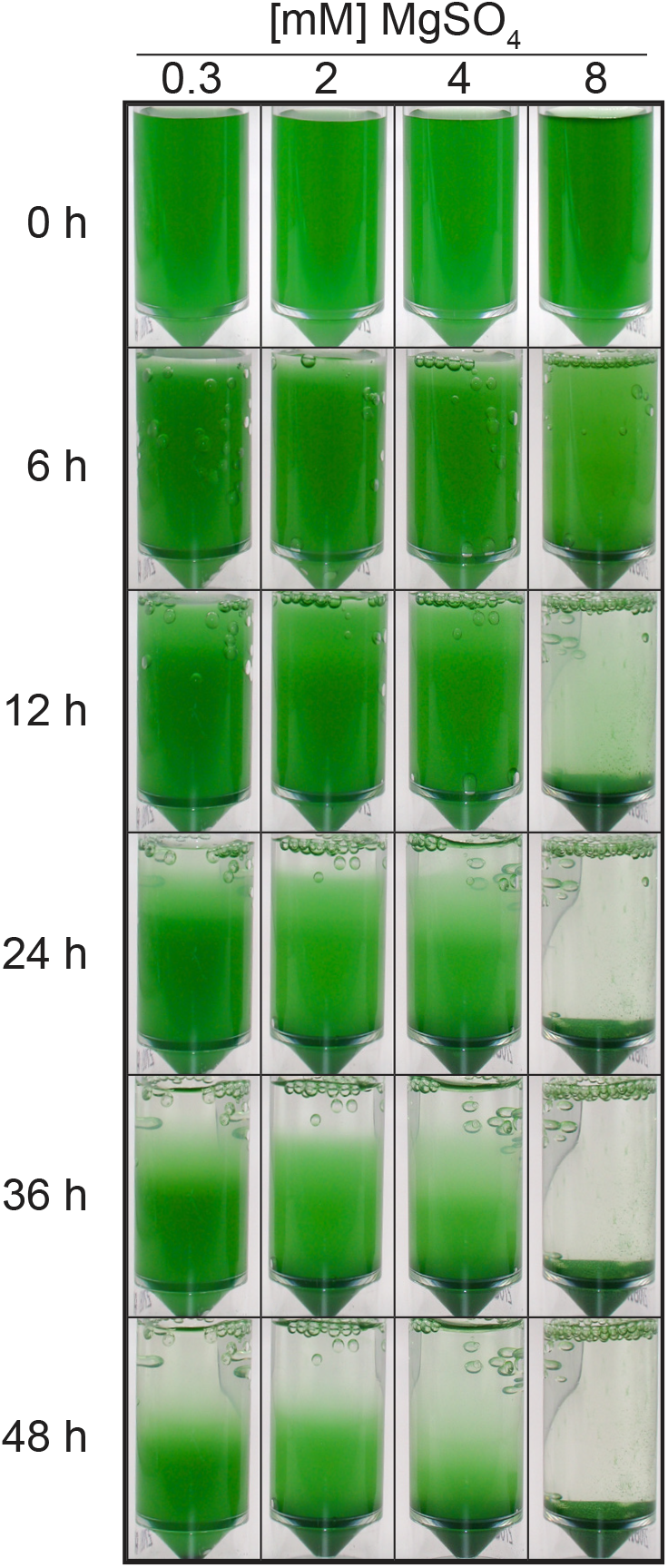
Sedimentation of 7942–KU (15 mL) cultivated in BG-11 with 0.3, 2, 4 and 8 mM MgSO_4._

However, the 24-hour sedimentation assay in universal tubes is not quantitative and does not easily allow for multiple testing. Therefore, to increase throughput and allow a quantitative comparison of variables, a miniaturised microplate-based sedimentation assay was developed. This assay relies on the observation that, in 48-well microplates, cells rapidly sediment to the centre of the well during static incubation (Figure 3A). A multipoint kinetic reading of the well absorbance then allows for an estimation of the relative amount of biomass that has accumulated in the centre of well relative to the initial absorbance reading. Thus, enabling the use of absorbance as a proxy for sedimentation. As a proof of concept, 7942–WT and 7942–KU were grown in BG-11 and P4 and multipoint absorbances were measured over a period of 20 minutes (Figure 3B). The accelerated sedimentation of 7942–KU was already visible at 5 min and continuously increased over the course of the measurement. Meanwhile, the sedimentation of the remaining strains was negligible throughout the whole period. The data from the microplate assay agree with what was previously observed in the 24-hour assay and offered a way to rapidly, and quantitatively, measure strain sedimentation.

**Figure 3.**
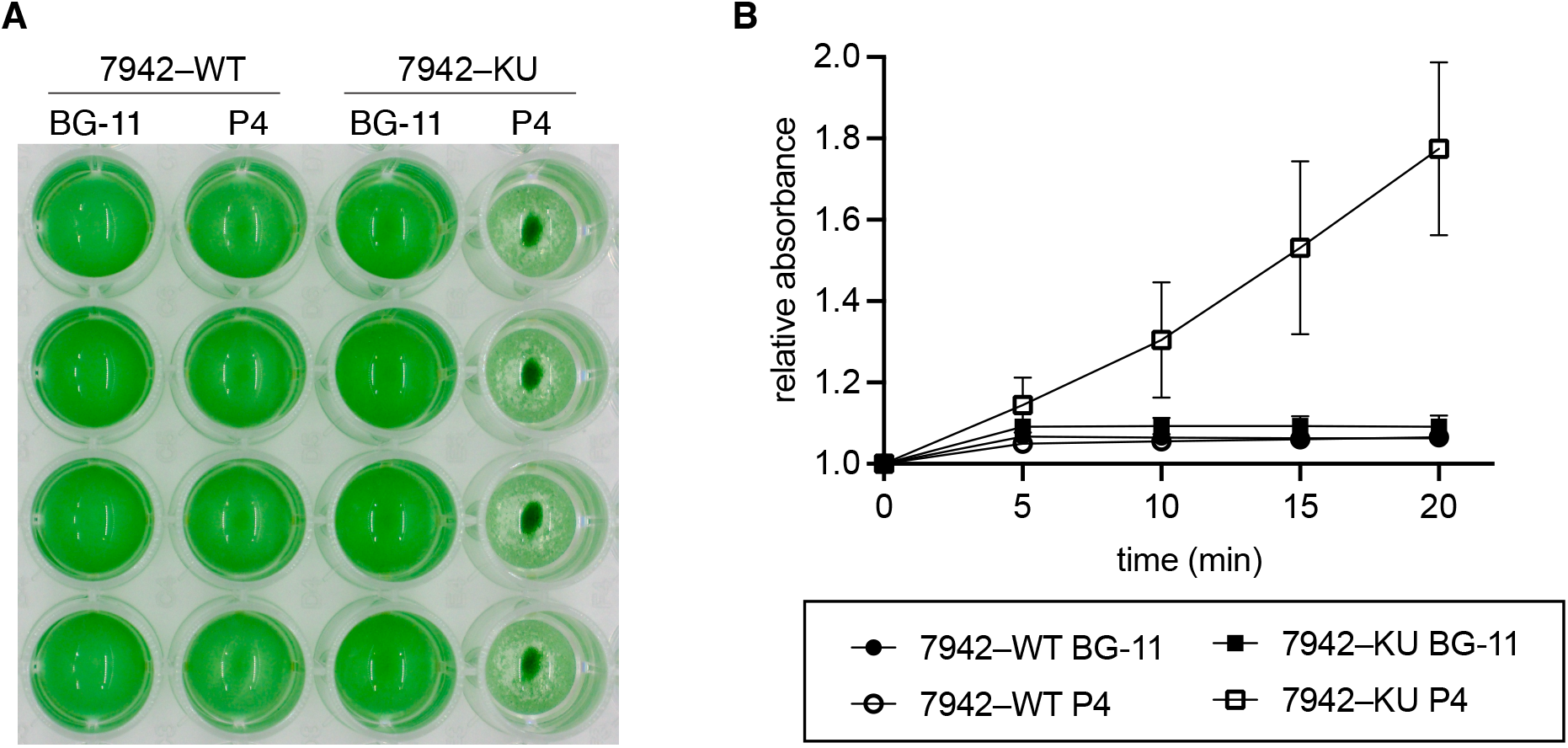
Development and testing of a high-throughput sedimentation assay. **A** Sedimentation of 7942–KU in a 48-well microplate after a 20 min static incubation. Four replicates of each sample are visible on the figure. **B** Microplate quantification of 7942– WT and 7942–KU sedimentation after cultivation in BG-11 and P4 media. n=4, error bars: standard deviation.

The microplate assay was then used to compare the effect of MgSO_4_, Na_2_SO_4_ and NaCl supplementation on sedimentation rates. For these experiments, 7942–WT and 7942– KU were grown in BG-11 medium to an exponential growth phase, then different salts were added to the cultures and sedimentation was measured after two hours. When the salts were added in the same concentration (12.5 mM), MgSO_4_ and Na_2_SO_4_ induced sedimentation, however there was no significant difference between the addition of NaCl and the control (Figure 4A). Considering the ionic strengths, 12.5 mM solutions of MgSO_4_, Na_2_SO_4_ and NaCl have ionic strengths of 50, 37.5 and 12.5 mM, respectively. Therefore, we tested adding the different salts in different concentrations to achieve the same ionic strength of 50 mM across all salts tested (Figure 4B). In this case, we observed sedimentation in all three salt treatments with no significant difference between them.

**Figure 4.**
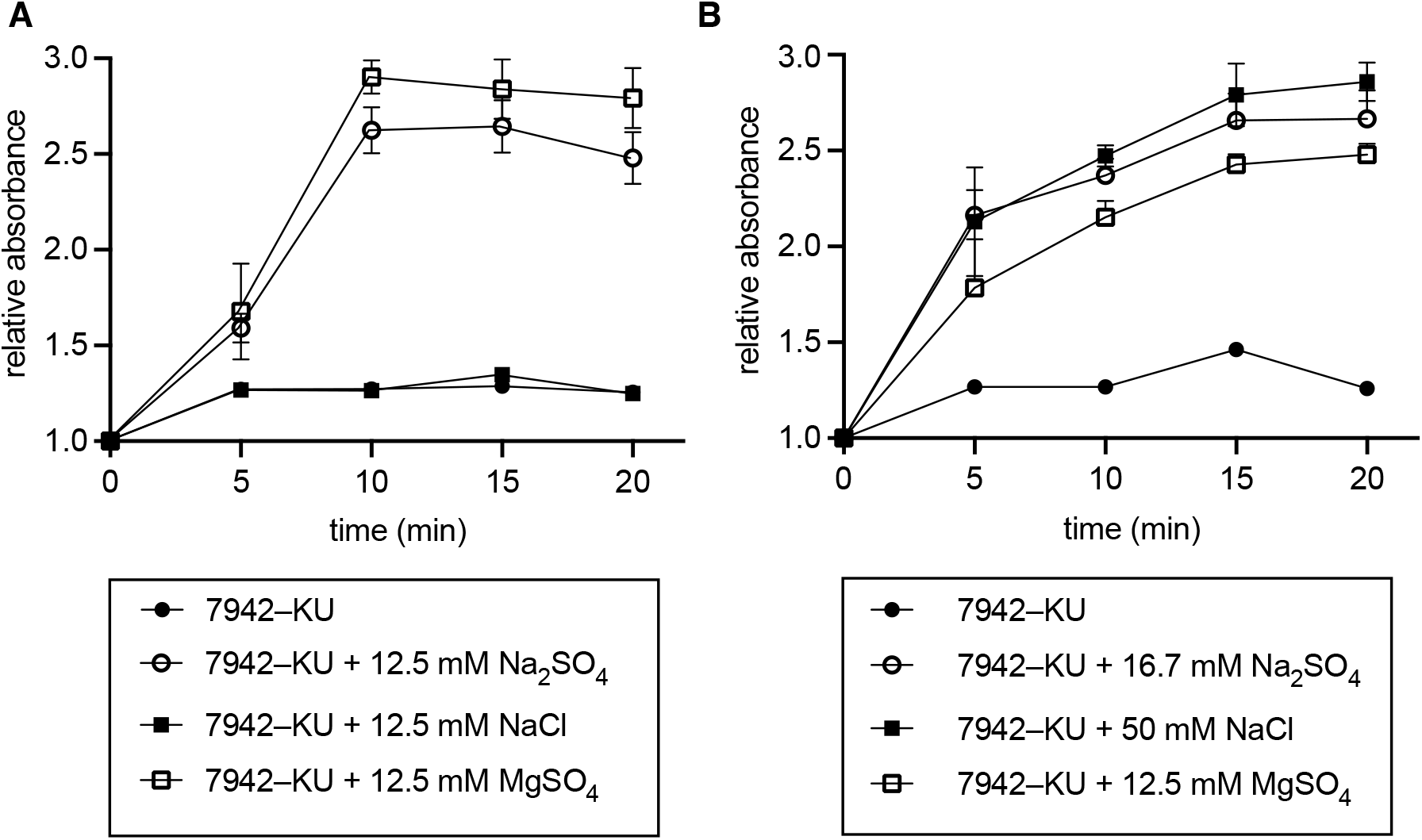
Microplate quantification of 7942–KU sedimentation after cultivation in BG-11 medium and addition of different salts. **A** 7942–KU sedimentation after addition of 12.5 mM of NaCl, Na_2_SO_4_ and MgSO_4_. **B** 7942–KU sedimentation after addition of 50, 16.7 and 12.5 mM of NaCl, Na_2_SO_4_ and MgSO_4_, respectively. n = 4, error bars: standard deviation.

### Unique mutations shed light on the history of S. elongatus 7942–KU

Having concluded that the rapid sedimentation was triggered by an increase in ionic strength of the medium, we aimed to determine the genetic differences that underlie this phenotype. 7942–KU was resequenced and, in comparison to the WT sequence (GenBank accession no. NC_007604), it has 8 SNPs in coding regions (Table 1) and 95 SNPs in non-coding regions. However, a recent study has shown that the 7942–WT reference sequence NC_007604 derives from a rare mutant clone and does not represent the strain typically used in the Golden lab, *S. elongatus* PCC 7942–AMC06 (hereafter 7942–AMC06) (Adomako *et al*., 2022). When comparing the 7942–KU coding SNPs with the 7942– AMC06 genome, it becomes clear that first, 7942–KU is closely related to 7942–AMC06 and second, out of the eight coding SNPs initially identified in 7942–KU, three also exist in 7942–AMC06 and another two are present in the same genes but encode different mutations (Table 1).

**Table 1.** 7942–KU and 7942–AMC06 SNPs in coding regions of the genome.

| 7942-WT locus | Locus description | 7942-KU SNPs | 7942-AMC06 SNPs (Adomako *et al.*, 2022) |
| --- | --- | --- | --- |
| Synpcc7942_0095 | Transcriptional regulator RpaA | Q121R | Q121R |
| Synpcc7942_0918 | Long-chain acyl-CoA synthetase Aas | L295P | L295P |
| Synpcc7942_0918 | Long-chain acyl-CoA synthetase Aas | T625I | T625I |
| Synpcc7942_1295 | Acyltransferase | del TCATGCC | S305I |
| Synpcc7942_1510 | Group 3 RNA polymerase sigma factor SigF | S239P | – |
| Synpcc7942_2070 | Type IVa pilus retraction ATPase pilT | C73Y | – |
| Synpcc7942_2373 | L-alanine-DL-glutamate epimerase | G264D | G264V |
| Synpcc7942_2597 | Adenylate cyclase | R89L | – |

The three 7942–KU genes with unique SNPs are *sigF* (Synpcc7942_1510), *pilT1* (Synpcc7942_2070) and an adenylate cyclase (Synpcc7942_2597). Given that a sedimentation phenotype has not been reported for 7942–AMC06, the three genes with unique mutations were further analysed. Synpcc7942_2070 encodes the type IVa pili (T4aP) retraction ATPase PilT1. Inactivation of PilT1 leads to the display of an excessive amount of pili on the cell surface (Bhaya *et al*., 2000). Hyperpiliated phenotypes can increase cell aggregation in *Synechocystis* (Conradi *et al*., 2019), however there are no reports linking hyperpiliation with an increase in sedimentation rate. Therefore, the subsequent work focused on *sigF* (Synpcc7942_1510) and the adenylate cyclase (Synpcc7942_2597). Synpcc7942_2597 encodes a putative adenylate cyclase. The mutation present in 7942–KU affects the conserved arginine at position 89 which completely abolishes enzyme activity (Keppetipola *et al*., 2007). It has been shown that phenotypes derived from the lack of intracellular cAMP can be complemented by the addition of exogenous cAMP (Terauchi and Ohmori, 1999; Bhaya *et al*., 2006). Therefore, to determine if the adenylate cyclase Synpcc7942_2597 could be responsible for the 7942–KU phenotype, cultures were grown in P4 medium, and the sedimentation assay was performed in the presence and absence of cAMP. No significant difference in sedimentation was observed (Supporting Information 1, Figure S1).

### Alterations in SigF and cell surface structures underpin fast sedimentation

Having ruled out the effect of cAMP on the sedimentation phenotype, we proceeded to investigate the role of *sigF* (Synpcc7942_1510) in the fast sedimentation phenotype. Synpcc7942_1510 encodes the group 3 RNA polymerase sigma factor SigF. The S239P mutation eliminates one of the conserved amino acids required for DNA binding and recognition of the –35 promoter element (Bhaya *et al*., 1999). Therefore, a 7942–WT SigF knockout mutant was generated and the sedimentation rate of the Δ*sigF* 7942–WT mutant was measured (Figure 5A, B). The Δ*sigF* mutant sedimented significantly faster than its WT counterpart. When compared to 7942–KU, this single mutation accounted for approximately 52% of the observed sedimentation. This result confirmed that SigF plays a significant role in the fast sedimentation phenotype, however it did not clarify the mechanism behind the fast sedimentation. In *Synechocystis*, SigF regulates polysaccharide release, vesiculation and cell surface composition (e.g. presence of T4aP) (Bhaya *et al*., 1999; Flores *et al*., 2019). Therefore, to determine if the fast sedimentation phenotype was a result of cell surface alterations, we took two approaches: measurement of the cell hydrophobicity and sedimentation analysis of an unpiliated mutant. First, a microbial adherence to hydrocarbons (MATH) assay (Rosenberg *et al*., 1980; Rosenberg, 1984) was performed with the three strains. This assay uses the level of bacterial adherence to an organic phase (typically an alkane) as a measure of the hydrophobicity of the cell surface. A more hydrophobic cell surface will result in a higher MATH score. We observed that both 7942–KU and the 7942–WT Δ*sigF* mutant had significantly less hydrophobic surfaces than 7942–WT (one-way ANOVA (F (2,9) = 30.88, p < .0001)) (Figure 6A). To determine if cell piliation was also a factor in the fast sedimentation phenotype, a 7942–WT Δ*pilB1* (Synpcc7942_2071) mutant was generated. PilB1 is the T4aP retraction ATPase, and its inactivation abrogates pili formation (Nagar *et al*., 2017). The sedimentation rate of 7942–WT Δ*pilB1* was measured and the results show that the strain sediments significantly faster than its WT counterpart and at virtually an identical rate to the Δ*sigF* mutant (Figure 6B). This result suggests that differences in the cyanobacterial cell surface account for the differential sedimentation observed between strains.

**Figure 5.**
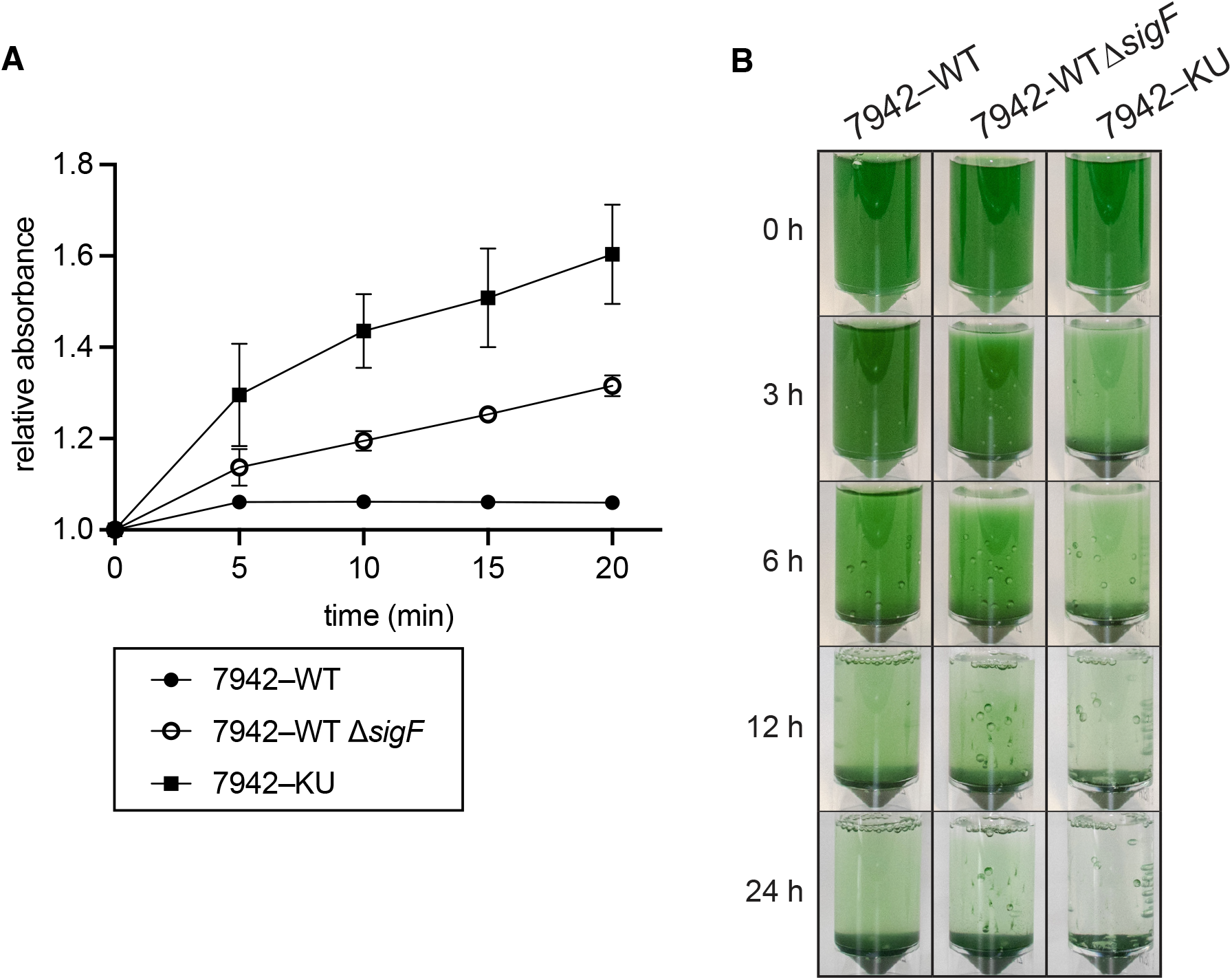
Sedimentation analysis of 7942–WT Δ*sigF* after cultivation in P4 medium. **A** Microplate quantification of 7942–WT, 7942–KU and 7942–WT Δ*sigF* sedimentation. n = 4, error bars: standard deviation. **B** 24-hour sedimentation assay (15 mL) in universal containers incubated under static conditions.

**Figure 6.**
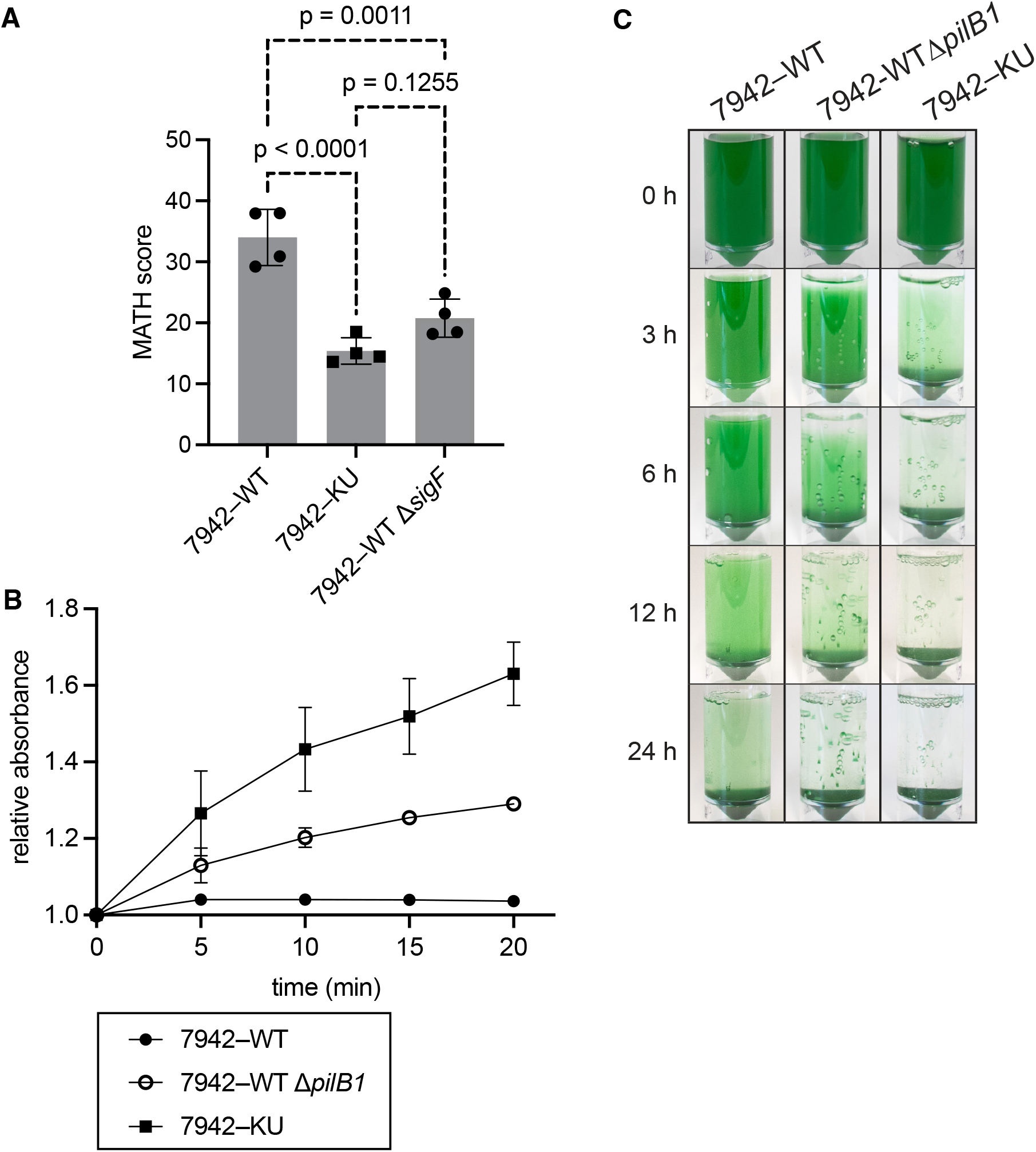
Cell surface hydrophobicity of 7942–WT, 7942–KU and 7942–WT Δ*sigF* and sedimentation analysis of 7942–WT Δ*pilB1*. **A** Surface hydrophobicity scores for 7942– WT, 7942–KU and 7942–WT Δ*sigF*. The microbial adhesion to hydrocarbons (MATH) assay was performed with cells cultivated in P4 medium. n = 4, error bars: standard deviation. P-values were determined using a one-way ANOVA followed by a Tukey’s multiple comparison test. **B** Microplate quantification of 7942–WT, 7942–KU and 7942–WT Δ*pilB1* sedimentation after cultivation in P4 medium. n = 4, error bars: standard deviation. **C** 24-hour sedimentation assay in universal containers incubated under static conditions.

## 3. Discussion

In this study we describe 7942–KU, a novel *S. elongatus* substrain that, through the process of laboratory domestication, has acquired a fast-sedimenting phenotype. We further demonstrate that the increased sedimentation is mediated by interactions between higher ionic strengths and altered cell surface characteristics of 7942–KU. Laboratory domestication is typically an undesired outcome of serial culturing. However, here we show that a detailed characterisation of domesticated cyanobacterial strains can provide insights into key cellular processes and open new perspectives for the biotechnological application of cyanobacteria.

### The cyanobacterial cell surface plays a significant role in the transition from a planktonic to a benthic state

The properties of the cyanobacterial cell surface have repeatedly been associated with regulating repulsion or attraction of cells. The mechanisms vary but typically include changes in cell surface structures (e.g., pili, S-layer), vesiculation and the composition of the extracellular matrix (Schatz *et al*., 2013; Allen *et al*., 2019; Conradi *et al*., 2019; Flores *et al*., 2019; Kera *et al*., 2020). In agreement with this, our findings suggest the 7942–KU fast sedimentation phenotype is caused by an altered cell surface. Cell surface modifications can increase the exposure of the cell to environmental fluctuations, thus leading to the observed fast sedimentation under higher ionic strengths. In the MATH assay, 7942–KU showed a significantly lower cell surface hydrophobicity (Figure 6A). This is in line with previous examples of increased cyanobacterial sedimentation when the cell surface hydrophobicity is decreased, or there is an increase in the hydrophobicity of the extracellular environment (Kawano *et al*., 2011; Jittawuttipoka *et al*., 2013; Li *et al*., 2021; Long *et al*., 2022).

As demonstrated by the 7942–WT mutants, a large part of the fast sedimentation can be traced back to deregulation of SigF and the loss of T4aP (Figs. 4, 5). However, it remains unclear whether SigF directly regulates T4aP expression or if this is the outcome of a pleiotropic effect of SigF on cell surface composition. In *Synechocystis*, SigF is associated with the regulation of cell surface dependent phenomena and it is thought to play a role in mediating reactions to salt and light stress (Bhaya *et al*., 1999; Huckauf *et al*., 2000). In addition, a *Synechocystis* Δ*sigF* mutant was described to exhibit cell envelope alterations such as impaired production of T4aP and outer membrane vesicles and a clumping phenotype (Flores *et al*., 2019). Therefore, it is not unlikely that SigF would fulfil similar roles in *S. elongatus*. However, neither the inactivation of SigF nor the loss of pili accounted completely for the level of sedimentation observed in 7942–KU. One possibility that could further contribute to the stronger sedimentation of 7942–KU is a change in exopolysaccharide (EPS) composition (both capsular and released). Several studies have shown that modulating the amount and composition of EPSs can drastically impact the repulsive forces between cells and their environment and lead to sedimentation (Allen *et al*., 2019; Conradi *et al*., 2019; Li *et al*., 2021). However, SNPs in genes involved in exopolysaccharide synthesis or release were not identified in the 7942–KU genome. Therefore, further investigation is required.

### Fast sedimenting strains are valuable for algal biotechnology

During microalgal production, harvesting can account for up to 30% of the total costs (Barros *et al*., 2015). Therefore, there is a keen interest in developing low-cost harvesting processes. Harvesting typically consists of a two-step process where cultures are concentrated by flocculation/sedimentation prior to employing higher cost physical cell separation techniques such as centrifugation or filtration. Many methods for large scale cell concentration have been proposed, however, they typically involve the addition of a flocculating agent or the application of an electric field (Barros *et al*., 2015). In the case of 7942–KU, full sedimentation can be achieved quickly (< 12 hours) without the addition of flocculating agents (Figs. 1, 2), thus allowing for easy product harvesting and recycling of the culture medium. The biotechnological use of self-aggregating cyanobacterial strains with modified cell surface properties has already been proposed. However, the previous studies either required long incubation times (> 15 days) or the secretion of a hydrophobic product (e.g., limonene) (Jittawuttipoka *et al*., 2013; Long *et al*., 2022). Conversely, 7942– KU requires only cultivation in a medium with a high ionic strength and is not tied to the production and secretion of hydrophobic compounds.

## 4. Concluding remarks

Laboratory domestication is a global phenomenon and virtually all model organisms have undergone this process. However, the impact of domestication on research outcomes is still underestimated. Going forward, it is important that genomic differences among substrains be considered during experimental planning. In addition, detailed investigation of domesticated substrains can help to elucidate how cyanobacteria adapt to a changing environment.

## 5. Materials and Methods

### Strains and growth conditions

In this study two strains of *Synechococcus elongatus* were used: *S. elongatus* PCC 7942 (obtained from the Pasteur Culture Collection of Cyanobacteria), here referred to as 7942– WT, and *Synechococcus elongatus* PCC 7942–KU (kindly donated by Yumiko Sakuragi, University of Copenhagen). Cultures were maintained on BG-11 medium (Stanier *et al*., 1971) supplemented with 10 mM 2-*tris*(hydroxymethyl)-methyl-2-amino 1-ethanesulfonic acid (TES) buffer (pH 8.0) and 1.5 % (*w*/*v*) Kobe I agar at 30°C with continuous illumination of approximately 25 μmol photons m^-1^ s^-1^. To prevent genetic drift, cultures were taken out of their respective cryo archives at least every three weeks. Liquid cultures were grown either in BG-11 TES medium or P4-TES CPH medium (Russo *et al*., 2019), a modified version of phosphate-replete medium (Lippi *et al*., 2018). Experiments in liquid culture were performed either in 100 mL flasks, T-25 cell culture flasks, 48-well flat-bottom microplates, or a CellDEG high-density system (HDC 6.10 starter kit; CellDeg, Germany). The CellDEG system was set up as previously described (Cao *et al*., 2022). All liquid cultures were illuminated by Lumilux cool white L 15 W/840 fluorescent lamps (Osram, Germany), with continuous illumination of approximately 50 μmol photons m^-2^ s^-1^, and shaken at 150 rpm (flasks, microplates) or 280 rpm (CellDEG, Germany) on a Unimax 1010 orbital shaker (Heidolph Instruments, Germany). For cultivation of the 7942–WT mutants, agar plates were supplemented with 25 μg mL^-1^ kanamycin (7942–WT Δ*pilB1*) or 5 μg mL^-1^ gentamycin (7942–WT Δ*sigF*). Antibiotics were not added to liquid cultures to avoid distorting effects.

### Sedimentation assays

For the sedimentation analysis in 100 mL flasks, cultures were grown for five days at 30°C under shaking and constant illumination. The cultures were then diluted to an OD_750 nm_ = 1 in 25 mL of the respective medium, transferred to 100 mL flasks and kept at room temperature, without agitation, under constant illumination, for approximately 14 days. For sedimentation analysis in 30 mL universal tubes and 48-well microplates, cultures were inoculated at an OD_750 nm_ = 0.4 and grown for 3 days at 30°C in the CellDEG system under shaking and constant illumination. For the assay in universal tubes, cultures were then diluted to an OD_750_ = 1.5 in 15 mL of the respective medium and kept at room temperature, without agitation, under constant illumination, for 24 to 48 h. For the microplate assay, cultures were diluted to an OD_750 nm_ = 1.0 in 200 μL of the respective medium and monitored with multipoint absorbance measurements (13 coordinates, OD_750 nm_) over 20 min on a Varioskan Flash microplate reader (Thermo Fisher Scientific, Germany). Microplates were kept in the reader at room temperature, without agitation or illumination during the assay. In the experiments where the addition of MgSO_4_, Na_2_SO_4_, NaCl or cAMP was tested, these components were added 2 h before the sedimentation analysis. The relative absorbance represents the ratio between the maximum value amongst the 13 coordinates at T_n_ and the mean of all 13 coordinates at T_0_ (relative absorbance = max T_n_/mean T_0_).

### Resequencing of S. elongatus 7942–KU

For sequencing 7942–KU, genomic DNA was extracted with the Quick-DNA Fungal/Bacterial Miniprep Kit (Zymo Research Europe, Germany) following the manufacturer’s recommendations with the following modifications. Cell lysis was conducted with Genomic Lysis Buffer in ZR BashingBead Lysis Tubes with a Vortex-Genie 2 (Scientific Industries, USA) on setting 7 for 20 min. The sample was then centrifuged at 10 000 x g for 2 min and the supernatant was column purified. Elution was performed in 100 μL of DNA Elution Buffer. To remove RNA from the sample, the eluate was brought to a final volume of 200 μL with DNA Elution Buffer and 10 μL of a 10 mg mL^-1^ solution of RNase A. The sample was gently mixed and incubated at 37°C for 60 min. A final clean-up was then conducted using the same column purification protocol. Genomic DNA was eluted in 50 μL of DNA Elution Buffer and yields were measured on a Qubit fluorometer (Invitrogen, Germany). Illumina 2 x 150 bp paired-end library preparation and sequencing was performed by Eurofins Genomics using their INVIEW Resequencing service for bacterial genomes and mapped against the 7942–WT genome (GenBank accession no. NC_007604). All SNPs identified in coding regions were confirmed by PCR and Sanger sequencing using the primers listed in Supporting Information 1, Table S1.

### Generation of S. elongatus PCC 7942 mutants

Constructs for the generation of knockout mutants were assembled by Gibson assembly (Gibson *et al*., 2009) using NEBuilder Hifi DNA Assembly Master Mix (New England Biolabs, USA). To generate a plasmid for the knockout of Synpcc7942_1510 (SigF), four DNA fragments were generated by PCR for plasmid assembly: a vector backbone of pBR322 without the tetracycline resistance cassette, 1000 bp genomic regions up- and downstream of Synpcc7942_1510 for homologous recombination and a gentamycin resistance cassette. The generated plasmid was named pJZ170. A plasmid for knocking out Synpcc7942_2071 (PilB1) was generated using a vector backbone fragment generated from pCAT.015 (Vasudevan *et al*., 2019) (gift from Alistair McCormick, Addgene plasmid # 119558) digested with XbaI and three fragments amplified by PCR: a Kanamycin resistance cassette and 1500 bp genomic regions up- and downstream of Synpcc7942_2071 for homologous recombination. This plasmid was named pMM02. Correct assembly of constructs was confirmed by Sanger sequencing. The generated constructs were transformed into 7942–WT using natural competence. Transformants were streaked on increasing concentrations of antibiotics (up to 10 μg mL^−1^ gentamycin and 50 μg mL^−1^ kanamycin respectively) until full segregation was confirmed by colony PCR. Fully segregated mutant strains were named 7942–WT Δ*sigF* (transformed with pJZ170) and 7942-WT Δp*ilB1* (transformed with pMM02). All primers used for generation of constructs and colony PCR are listed in Supporting Information 1, Table S2.

### Microbial adhesion to hydrocarbons (MATH) assay

The MATH assay was performed following the original publication (Rosenberg *et al*., 1980) with some modifications. Exponentially growing cyanobacterial cells were harvested and washed twice in phosphate buffer (22.2 g L^-1^ K_2_HPO_4_, 7.2 g L^-1^ KH_2_PO_4_), pH 7.1. Following this, cells were resuspended to, approximately, an OD_750 nm_ = 0.35 in 1.6 mL of phosphate buffer with 2 M ammonium sulphate in glass vials. The addition of ammonium sulphate enhances hydrophobic interactions and, thus, the sensitivity of the assay (Rosenberg, 1984). To the 1.6 mL cyanobacterial suspension, 400 μL of *p*-xylene were added, the mixture was vortexed at maximum speed for 2 min and then allowed to rest for 15 min. 800 μL were harvested from the aqueous phase and the OD_750 nm_ was measured. The MATH score is calculated using the equation (1 − OD_final_/OD_initial_) × 100.

## 6. Acknowledgements

The authors thank Yumiko Sakuragi for providing the *S. elongatus* PCC 7942–KU strain. DAR was supported by the Alexander von Humboldt Foundation and by the IMPULSE^project^ (IP-2020-03, Friedrich Schiller University Jena). This work was funded by the Deutsche Forschungsgemeinschaft (DFG, German Research Foundation) under Germany’s Excellence Strategy – EXC 2051 – Project-ID 390713860 (JAZZ) and by the Deutsche Forschungsgemeinschaft (DFG, German Research Foundation), SFB 1127 ChemBioSys, project number 239748522 (JAZZ).

## 8. Author contributions

Conceptualization: DAR; Methodology: JAZZ, MM, DAR; Investigation: JAZZ, MM, DAR; Resources: JAZZ, DAR, GP; Data curation: JAZZ, MM, DAR; Formal analysis: JAZZ, MM, DAR; Visualization: JAZZ, MM, DAR; Writing – original draft: DAR, JAZZ; Writing – review & editing: JAZZ, MM, GP, DAR; Funding acquisition: DAR. All authors revised the manuscript and approved its final version.

## 9. Data availability statement

Raw read files and variant analysis from the Eurofins Genomics resequencing of PCC 7942–KU can be found at 10.5281/zenodo.7299545. Δ*sigF* and Δ*pilB1* plasmid maps can be found at 10.5281/zenodo.7299545. PCC 7942–KU is available from the DSMZ – German Collection of Microorganisms and Cell Cultures (DSM no. XXXXX).

## References

Adomako, M., Ernst, D., Simkovsky, R., Chao, Y.-Y., Wang, J., Fang, M., et al. (2022) Comparative Genomics of Synechococcus elongatus Explains the Phenotypic Diversity of the Strains. mBio 13: e00862–22.

Allen, R., Rittmann, B.E., and Curtiss, R. (2019) Axenic Biofilm Formation and Aggregation by Synechocystis sp. Strain PCC 6803 Are Induced by Changes in Nutrient Concentration and Require Cell Surface Structures. Appl Environ Microbiol 85: e02192–18.

Barros, A.I., Gonçalves, A.L., Simões, M., and Pires, J.C.M. (2015) Harvesting techniques applied to microalgae: A review. Renewable and Sustainable Energy Reviews 41: 1489–1500.

Bhaya, D., Bianco, N.R., Bryant, D., and Grossman, A. (2000) Type IV pilus biogenesis and motility in the cyanobacterium Synechocystis sp. PCC 6803. Molecular Microbiology 37: 941–951.

Bhaya, D., Nakasugi, K., Fazeli, F., and Burriesci, M.S. (2006) Phototaxis and Impaired Motility in Adenylyl Cyclase and Cyclase Receptor Protein Mutants of Synechocystis sp. Strain PCC 6803. Journal of Bacteriology 188: 7306–7310.

Bhaya, D., Watanabe, N., Ogawa, T., and Grossman, A.R. (1999) The role of an alternative sigma factor in motility and pilus formation in the cyanobacterium Synechocystis sp. strain PCC6803. PNAS 96: 3188–3193.

Bryant, D.A. (2014) A Brief History of Cyanobacterial Research: Past, Present, and Future Prospects. In The Cell Biology of Cyanobacteria. Caister Academic Press, pp. 1–5.

Cao, J., Russo, D.A., Xie, T., Groß, G.A., and Zedler, J.A.Z. (2022) A droplet-based microfluidic platform enables high-throughput combinatorial optimization of cyanobacterial cultivation. Sci Rep 12: 15536.

Cohen, S.E. and Golden, S.S. (2015) Circadian Rhythms in Cyanobacteria. Microbiol Mol Biol Rev 79: 373–385.

Conradi, F.D., Zhou, R.-Q., Oeser, S., Schuergers, N., Wilde, A., and Mullineaux, C.W. (2019) Factors Controlling Floc Formation and Structure in the Cyanobacterium Synechocystis sp. Strain PCC 6803. Journal of Bacteriology 201: e00344–19.

Eydallin, G., Ryall, B., Maharjan, R., and Ferenci, T. (2014) The nature of laboratory domestication changes in freshly isolated Escherichia coli strains. Environmental Microbiology 16: 813–828.

Flores, C., Santos, M., Pereira, S.B., Mota, R., Rossi, F., De Philippis, R., et al. (2019) The alternative sigma factor SigF is a key player in the control of secretion mechanisms in Synechocystis sp. PCC 6803. Environmental Microbiology 21: 343–359.

Gibson, D.G., Young, L., Chuang, R.-Y., Venter, J.C., Hutchison, C.A., and Smith, H.O. (2009) Enzymatic assembly of DNA molecules up to several hundred kilobases. Nat Methods 6: 343–345.

Golden, S.S. (2019) The international journeys and aliases of Synechococcus elongatus. New Zealand Journal of Botany 57: 70–75.

Huckauf, J., Nomura, C., Forchhammer, K., and Hagemann, M.Y. 2000 (2000) Stress responses of Synechocystis sp. strain PCC 6803 mutants impaired in genes encoding putative alternative sigma factors. Microbiology 146: 2877–2889.

Ikeuchi, M. and Tabata, S. (2001) Synechocystis sp. PCC 6803 — a useful tool in the study of the genetics of cyanobacteria. Photosynthesis Research 70: 73–83.

Jittawuttipoka, T., Planchon, M., Spalla, O., Benzerara, K., Guyot, F., Cassier-Chauvat, C., and Chauvat, F. (2013) Multidisciplinary Evidences that Synechocystis PCC6803 Exopolysaccharides Operate in Cell Sedimentation and Protection against Salt and Metal Stresses. PLOS ONE 8: e55564.

Kaneko, T., Sato, S., Kotani, H., Tanaka, A., Asamizu, E., Nakamura, Y., et al. (1996) Sequence analysis of the genome of the unicellular cyanobacterium Synechocystis sp. strain PCC6803. II. Sequence determination of the entire genome and assignment of potential protein-coding regions. DNA Res 3: 109– 136.

Kawano, Y., Saotome, T., Ochiai, Y., Katayama, M., Narikawa, R., and Ikeuchi, M. (2011) Cellulose Accumulation and a Cellulose Synthase Gene are Responsible for Cell Aggregation in the Cyanobacterium Thermosynechococcus vulcanus RKN. Plant and Cell Physiology 52: 957–966.

Keppetipola, N., Jain, R., and Shuman, S. (2007) Novel Triphosphate Phosphohydrolase Activity of Clostridium thermocellum TTM, a Member of the Triphosphate Tunnel Metalloenzyme Superfamily*. Journal of Biological Chemistry 282: 11941– 11949.

Kera, K., Yoshizawa, Y., Shigehara, T., Nagayama, T., Tsujii, M., Tochigi, S., and Uozumi, N. (2020) Hik36–Hik43 and Rre6 act as a two-component regulatory system to control cell aggregation in Synechocystis sp. PCC6803. Sci Rep 10: 19405.

Li, J., Fang, D., Ye, R., Zhou, C., and Li, P. (2021) The released polysaccharide inhibits cell aggregation and biofilm formation in the cyanobacterium Synechocystis sp. PCC 6803. European Journal of Phycology 56: 119–128.

Li, S., Sun, T., Xu, C., Chen, L., and Zhang, W. (2018) Development and optimization of genetic toolboxes for a fast-growing cyanobacterium Synechococcus elongatus UTEX 2973. Metabolic Engineering 48: 163–174.

Lippi, L., Bähr, L., Wüstenberg, A., Wilde, A., and Steuer, R. (2018) Exploring the potential of high-density cultivation of cyanobacteria for the production of cyanophycin. Algal Research 31: 363–366.

Long, B., Fischer, B., Zeng, Y., Amerigian, Z., Li, Q., Bryant, H., et al. (2022) Machine learning-informed and synthetic biology-enabled semi-continuous algal cultivation to unleash renewable fuel productivity. Nat Commun 13: 541.

Nagar, E., Zilberman, S., Sendersky, E., Simkovsky, R., Shimoni, E., Gershtein, D., et al. (2017) Type 4 pili are dispensable for biofilm development in the cyanobacterium Synechococcus elongatus. Environ Microbiol 19: 2862–2872.

Rosenberg, M. (1984) Ammonium sulphate enhances adherence of Escherichia coli J-5 to hydrocarbon and polystyrene. FEMS Microbiology Letters 25: 41–45.

Rosenberg, M., Gutnick, D., and Rosenberg, E. (1980) Adherence of bacteria to hydrocarbons: A simple method for measuring cell-surface hydrophobicity. FEMS Microbiology Letters 9: 29–33.

Russo, D.A., Zedler, J.A.Z., Conradi, F.D., Schuergers, N., Jensen, P.E., Mullineaux, C.W., et al. (2022) Development of a Highly Sensitive Luciferase-Based Reporter System To Study Two-Step Protein Secretion in Cyanobacteria. J Bacteriol 204: e00504–21.

Russo, D.A., Zedler, J.A.Z., Wittmann, D.N., Möllers, B., Singh, R.K., Batth, T.S., et al. (2019) Expression and secretion of a lytic polysaccharide monooxygenase by a fast-growing cyanobacterium. Biotechnol Biofuels 12: 74.

Santos-Merino, M., Singh, A.K., and Ducat, D.C. (2019) New Applications of Synthetic Biology Tools for Cyanobacterial Metabolic Engineering. Frontiers in Bioengineering and Biotechnology 7:.

Schatz, D., Nagar, E., Sendersky, E., Parnasa, R., Zilberman, S., Carmeli, S., et al. (2013) Self-suppression of biofilm formation in the cyanobacterium Synechococcus elongatus. Environmental Microbiology 15: 1786–1794.

Schirmacher, A.M., Hanamghar, S.S., and Zedler, J.A.Z. (2020) Function and Benefits of Natural Competence in Cyanobacteria: From Ecology to Targeted Manipulation. Life 10: 249.

Schuergers, N., Lenn, T., Kampmann, R., Meissner, M.V., Esteves, T., Temerinac-Ott, M., et al. (2016) Cyanobacteria use micro-optics to sense light direction. eLife 5: e12620.

Stanier, R.Y., Kunisawa, R., Mandel, M., and Cohen-Bazire, G. (1971) Purification and properties of unicellular blue-green algae (order Chroococcales). Bacteriological Reviews 35: 171–205.

Taton, A., Erikson, C., Yang, Y., Rubin, B.E., Rifkin, S.A., Golden, J.W., and Golden, S.S. (2020) The circadian clock and darkness control natural competence in cyanobacteria. Nature Communications 11: 1688.

Terauchi, K. and Ohmori, M. (1999) An Adenylate Cyclase, Cyal, Regulates Cell Motility in the Cyanobacterium Synechocystis sp. PCC 6803. Plant Cell Physiol 40: 248– 251.

Trautmann, D., Voß, B., Wilde, A., Al-Babili, S., and Hess, W.R. (2012) Microevolution in Cyanobacteria: Re-sequencing a Motile Substrain of Synechocystis sp. PCC 6803. DNA Res 19: 435–448.

Ungerer, J., Lin, P.-C., Chen, H.-Y., and Pakrasi, H.B. (2018) Adjustments to Photosystem Stoichiometry and Electron Transfer Proteins Are Key to the Remarkably Fast Growth of the Cyanobacterium Synechococcus elongatus UTEX 2973. mBio 9: e02327–17.

Ungerer, J., Wendt, K.E., Hendry, J.I., Maranas, C.D., and Pakrasi, H.B. (2018) Comparative genomics reveals the molecular determinants of rapid growth of the cyanobacterium Synechococcus elongatus UTEX 2973. Proceedings of the National Academy of Sciences 115: E11761–E11770.

Vandamme, D., Foubert, I., Fraeye, I., Meesschaert, B., and Muylaert, K. (2012) Flocculation of Chlorella vulgaris induced by high pH: Role of magnesium and calcium and practical implications. Bioresource Technology 105: 114–119.

Vasudevan, R., Gale, G.A.R., Schiavon, A.A., Puzorjov, A., Malin, J., Gillespie, M.D., et al. (2019) CyanoGate: A Modular Cloning Suite for Engineering Cyanobacteria Based on the Plant MoClo Syntax. Plant Physiology 180: 39–55.

Williams, J.G.K. (1988) [85] Construction of specific mutations in photosystem II photosynthetic reaction center by genetic engineering methods in Synechocystis 6803. In Methods in Enzymology. Cyanobacteria. Academic Press, pp. 766–778.

Yang, Y., Lam, V., Adomako, M., Simkovsky, R., Jakob, A., Rockwell, N.C., et al. (2018) Phototaxis in a wild isolate of the cyanobacterium Synechococcus elongatus. Proceedings of the National Academy of Sciences 115: E12378–E12387.

Yu, J., Liberton, M., Cliften, P.F., Head, R.D., Jacobs, J.M., Smith, R.D., et al. (2015) Synechococcus elongatus UTEX 2973, a fast growing cyanobacterial chassis for biosynthesis using light and CO_2_. Scientific Reports 5: 8132.

