## Supporting Information 1 for "Cell surface composition and ionic strength mediate fast sedimentation in the cyanobacterium *Synechococcus elongatus* PCC 7942"

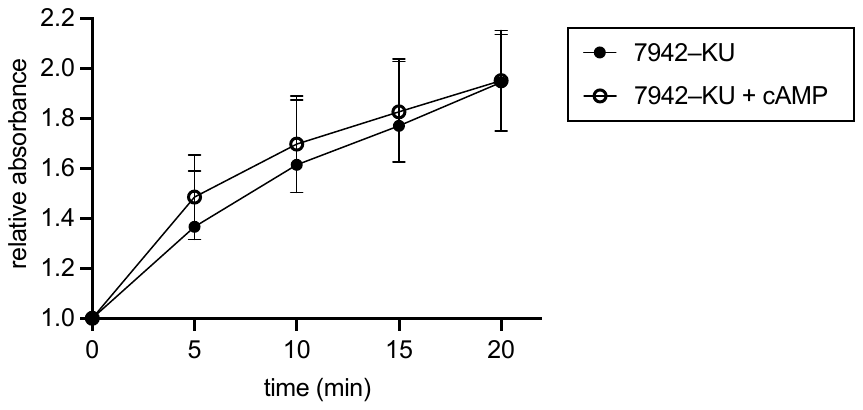

**Figure S1** Quantification of 7942–KU sedimentation after cultivation in P4 medium and static incubation in the presence and absence of 100 µM cAMP. n = 4, error bars: standard deviation.

**Table S1** Primers used for confirmation of SNPs in 7942–KU by PCR and Sanger sequencing.

| Primer name | Primer sequence (5’-3’) |
| --- | --- |
| SYNPCC7942_0095 F | CAG AAC AGA GCC ACT ATC ATT T |
| SYNPCC7942_0095 R | GAA CAG GCA CAG AAA CAA CT |
| SYNPCC7942_0918 F | CAA GTC ACA ACA TTA GGT GTG G |
| SYNPCC7942_0918 R | GAG TAC CGA GGG TCA TAG ATA |
| SYNPCC7942_1295 F | CCA AGT ATC AAT GGG CAT CGA |
| SYNPCC7942_1295 R | AAC GAA CTG GTG GAC GAT T |
| SYNPCC7942_1510 F | CAA AGC TTG TGC GAT TCT TA |
| SYNPCC7942_1510 R | TCG TAT TTT GTT TGC AGA CC |
| SYNPCC7942_2070 OE F | GTA GGA ATT CAT GGA GCT GAT GAT TGA AGA CC |
| SYNPCC7942_2070 OE R | TAT CAA GCT TCT ATG CAG CAA CCG GAA CAT TA |
| SYNPCC7942_2373 F | GCT TTG TTC ATT TGG CAG C |
| SYNPCC7942_2373 R | TCT CGT GAT CAA TCA CCG CT |
| SYNPCC7942_2597 F | GGT GAG CAG GCA ATT AAA GC |
| SYNPCC7942_2597 R | GAG ATG GCA ACC GTC AAA T |

**Table S2.** Primers used for generation of plasmids and knockout mutants.

| Primer name | Primer sequence (5’-3’) | Usage of primer |
| --- | --- | --- |
| SigF-Flank1-GibF | AAC GCA GTC AGG CAC CGT GTC AAG GCT ACG CCG TAG AAG GG | Amplification of Gibson fragments for the generation of pJZ170 (Synpcc7942_1510, SigF knockout vector) |
| SigF-Flank1-GibR | GTC CTG GCT GGC GAA CGA GCA CTG GCT CCG TTG TTC GCT |  |
| SigF-Flank2-GibF | GTA AAT TGT CAC AAC GCC GCA TTG CTA ACC AAG CAG AGG CAG |  |
| SigF-Flank2-GibR | GAG GTG CCG CCG GCT TCC ATC AGC AGC CGC GAT CTC AAT ATG |  |
| pBR-GibF | ATG GAA GCC GGC GGC ACC TC |  |
| pBR-GibR | ACA CGG TGC CTG ACT GCG TTA GC |  |
| GenGib-F | GCT CGT TCG CCA GCC AGG AC |  |
| GenGib-R | GCG GCG TTG TGA CAA TTT ACC GAA C |  |
| PilB1 downflank F | TGC TCG ATG AGT TTT TCT AAC AGT CAG CAG CGA TCG TGC A | Amplification of Gibson fragments for the generation of pMM02 (Synpcc7942_ 2071, PilB1 knockout vector) |
| PilB1 downflank R | TGC CTG CAG GTC GAC TCT AGG TTT TAC GAA GCT TGG GAT TAC GTG T |  |
| PilB1 upflank F | TTC GAA GAC AAG GCA TCT AGC GGC GAT CGC TCC AGC GTC C |  |
| PilB1 upflank R | TTG AGA CAC AAC GTG GCT TTG GGT GGC AGT GGC TGA GAG GAA AA |  |
| KanR amp F | AAA GCC ACG TTG TGT CTC AA |  |
| KanR amp R | TTA GAA AAA CTC ATC GAG CAT CAA |  |
| SigF_intR | ATT GCT TGC TGG AGT CGA ATC | Colony PCR to confirm segregation of 7942-WT∆SigF (Synpcc7942_1510 knockout) |
| SigF_FlankR | CTT CAT GTA GCA AAT GTG AAG TCG |  |
| SigF_FlankF | GCG AGA TTG TTG ATC AGA GTG |  |
| Genta R | TGC TTG CAC GTA GAT CAC |  |
| PilB1 KO seq F | AGA GAC TCC AGA AAG ACA GC | Colony PCR to confirm segregation of 7942-WT∆PilB1 (Synpcc7942_2071 knockout) |
| PilB1 colony PCR R | ACG AGA ATG AGA CCA TAG GG |  |
| KanR seq R | ACA AAC AGG AAT CGA ATG CA |  |

**Table S3** Composition of BG-11 and P4 media used in this study.

| Component | BG-11 TES | P4-CPH TES medium |
| --- | --- | --- |
| NaNO_3_ | 17.6 mM | 50 mM |
| KNO_3_ | – | 15 mM |
| MgSO_4_ ⋅7 H_2_O | 0.3 mM | 2 mM |
| CaCl_2_ ⋅ 2 H_2_O | 0.245 mM | 0.5 mM |
| K_2_HPO_4_ ⋅3 H_2_O | 0.175 mM | – |
| KH_2_PO_4_ | – | 4 mM |
| FeCl_3_ ⋅ 6 H_2_O | – | 150 µM |
| Ferric ammonium citrate | 0.6 mg/L | – |
| Citric acid | 3.1 µM | – |
| EDTA Na_2_ salt dihydrate | 2.7 µM | 150 µM |
| Na_2_CO_3_ | 188.7 µM | – |
| H_3_BO_3_ | 46.26 µM | 25 µM |
| MnCl_2_ ⋅ 4 H_2_O | 9.1 µM | 20 µM |
| ZnSO_4_ ⋅ 7 H_2_O | 0.8 µM | 2 µM |
| Na_2_MoO_4_ ⋅ 2 H_2_O | 1.6 µM | 3 µM |
| CuSO_4_ ⋅ 5 H_2_O | 0.3 µM | 0.02 µM |
| Co(NO_3_)_2_ ⋅ 6 H_2_O | 0.2 µM | – |
| CoCl_2_ ⋅ 6 H_2_O | – | 0.06 µM |
| C_6_H_15_NO_6_S (TES) | 10 mM | 10 mM |
